# Candidate gene prioritization using graph embedding

**DOI:** 10.1101/2020.02.03.927913

**Authors:** Quan Do, Pierre Larmande

## Abstract

Candidate genes prioritization allows to rank among a large number of genes, those that are strongly associated with a phenotype or a disease. Due to the important amount of data that needs to be integrate and analyse, gene-to-phenotype association is still a challenging task. In this paper, we evaluated a knowledge graph approach combined with embedding methods to overcome these challenges. We first introduced a dataset of rice genes created from several open-access databases. Then, we used the Translating Embedding model and Convolution Knowledge Base model, to vectorize gene information. Finally, we evaluated the results using link prediction performance and vectors representation using some unsupervised learning techniques.

## I. Introduction

Rice is the first global cereal in terms of production / human diet but also a model plant for agronomic research. In order to meet the challenges of global food growth in a context of climate change, especially in Vietnam who is a major producer, a better understanding of genotype-phenotype relationships is crucial to improve production capabilities. Recent advances in plant biotechnology lead to an unprecedented revolution in the acquisition of diverse data: phenotypic, transcriptomic, genomic, etc. However, data currently available is highly distributed and patchy in nature. For scientists, the challenge lies in integrating data and finding relevant information as the amount of data is hard to manage. For instance, Genome-wide association studies (GWAS) usually provide several hundred to several thousand genes potentially associated with a phenotype. Since using pure experimental approaches to verify all candidates could be expensive, a number of computational methods have been developed to rank candidates genes [1]–[4]. Although the number of deep learning applications in genomics is increasing, few of them handle the challenge of genes prioritization [5], [6]. In this paper, we applied graph embeddings techniques to study and rank gene-phenotype interactions.

### A. Databases

Currently, rice gene information is available in various online databases where each of them contain different attributes. The challenge is to collect comprehensive data and organise it logically to enable efficient analysis. In order to build a new dataset, we first considered several rice databases containing complementary information: Oryzabase [7], The Rice Annotation Project (RAP) [8], Rice SNP-Seek Database [9], Funricegene [10], The Universal Protein Resource (UniProt) [11], Gramene [12], Rice Expression Database (RED) [13], MSU Rice Genome Annotation Project (RGAP) [14], Predicted Rice Interactome Network (PRIN) [15]. Then, we used the PyRice package [16] to collect data and filter important information. More details of the dataset are available in section III-A.

### B. Knowledge graphs

Knowledge graphs (KGs) represent entity information and their relationship. The representation of knowledge has a long history of logic and artificial intelligence. Knowledge graphs are similar to simple graphs which include two components: Vertex/Entity and Edge/Relation. A knowledge graphs is a directional graph so we can represent it in term of the original form, including sets of triples. A triple includes head entity, relation, tail entity, denoted as (h, r, t), (e.g. BarackObama, wasBornIn, Honolulu). Knowledge graphs have wide of benefits, based on the definition of vertex and edge such as: semantic searching and ranking [17], [18], question answering [19] and machine reading [20].

The critical point of knowledge graphs is how to store graphs efficiently so that we can extract as much information as possible. Moreover the approach must be fast enough (and simple) to deploy in real-time systems. The Knowledge graph embedding approach allows to represent entities as points and relations as scalar vectors in space coordinates. However, in practical, knowledge graphs usually consist of millions or billions of triples, while many of them are invalid or missing [21]. Many studies are focusing on the task of improving knowledge graphs to predict the missing triples in KGs, such as predicting whether a triple not in KGs is likely to be valid or not [22].

Recently, many embedding models have been proposed to learn the vector representation of the components in the triples. The link prediction results, introduced by Nickel et al. [23] obtains the state-of-the-art. Embedding models evaluate the triple (h, r, t), optimally so that the valid triple are higher than the invalid triples.

Conventional embedding methods such as DISTMULT [24], TransE [25], ComplEx [26] have been successfully applied in the knowledge extraction fields. However, these approaches only aim to explore the linear relationship among entities because they are mostly based on simple operators such as addition, subtraction or multiplication. In the last decades, convolution neural networks, introduced by LeCun et al. [27], were initially designed for computer vision. Recently, based on the research of Collobert [28], [29], CNN received a significant attention in the field of natural language processing. By learning non-linear features to capture the relationships between components, CNN usually takes less number of parameters compared to standard fully-connected neural networks thereby reducing the computation cost. Inspired by the success of CNN, Dettmers et al. [30] proposed a model named ConvE -the first model applying CNN for the knowledge extraction task.

Moreover, ConvKB, introduced by Nguyen et al. [31], is a CNN based model for knowledge graph completion which successfully achieved state-of-the-art results. In practical, knowledge graph embedding models are commonly constructed to model entries at the same dimension for the given triple, where presumably each dimension captures some relation-specific attribute of entities [32]. However, following the research of [32], existing models are not available to provide a deep architecture to model the relationship among entities in triples on the same dimension.

### C. Objective

In the scope of this paper, our contributions are descibed as follows:

- Introduce a new dataset named *OsGenePrio* for rice gene information and pre-process the data to use the embedding models TransE and ConvKB in order to explore information in a KGs context.
- Evaluate the results of the embedding models using link prediction performance (Mean rank - MR, Mean reciprocal rank - MRR and Hit ratio - Hits@). After representing gene information as vectors, we used unsupervised learning techniques: to cluster genes (using K-Means Clustering), to find similarity between genes (using K-Nearest Neighbors).

## II. Embedding Model

A knowledge base 𝒢 represents a set of triples (*h, r, t*) with *h, t* ∈ *ε* and *r* ∈ ℛ, where *ε* is a set of entities and ℛ is a set of relations. Embedding models are used to define a score function *(score function) f*, which provides the score for each triple (*h, r, t*) such the score of the real triples is always higher than the unreal triples. For each triple (*h, r, t*), the corresponding vector *k* dimensions is marked as *v*_*h*_, *v*_*r*_, *v*_*t*_ which correspond to an input 3 vectors *k* dimension.

### A. Translating Embedding model

Translating Embedding (TransE) model is the most basic among several types of embedding models. The idea of TransE aims to optimize the sum of the head entity and relation as close as possible with the tail entity. A triple including head entity, relation, tail entity, denoted as *h, r, t*. In general, a triple is represented as:

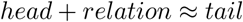

or with format vector *v*_*h*_, *v*_*r*_, *v*_*t*_:

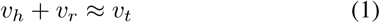

For example, a triple (BarackObama, wasBornIn, Honolulu) satisfy this property:

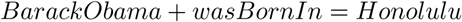

The original TransE has based on the reduce of loss function to improve model performance.

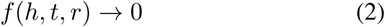

In practical, the most basic case of TransE is represented as:

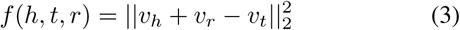

However, there is a drawback of TransE because it only covers the case of one-to-one relation, and is not suitable for one-to-many/many-to-one relation. For example, if there exists two triples with similar components such as (BarackObama, wasBornIn, Honolulu) and (BrunoMars, wasBornIn, Honolulu), the entity vector of “‘BarackObama’” will be very close to the other one after the training phase. Nevertheless, there is no such similarity in the real world. To handle this, we used the TransE model as pre-trained to replace the initial random entities vectors.

### B. Convolution Knowledge Base model

#### 1) Idea

In this project, we used an embedded Convolution Knowledge Base (ConvKB) model to complete a knowledge base. The ConvKB model is designed to capture the relationship between entities in the knowledge base.

In the ConvKB model, each entity or relation is represented as a *k* dimensional vector. For each triple (*h, r, t*), *v*_*h*_, *v*_*r*_, *v*_*t*_ which corresponds to an input matrix size *k*×3. This matrix is processed on the convolutional layer where each different filter having the same size of 1 × 3, is used to extract the general relation in a triple. These filters slide on rows of matrix to create different feature maps. Then, these feature maps are concatenate to form a feature vector and together with the weight vector, will return a score for the triple (*h, r, t*). This score is used to evaluate if the triple is valid (*h, r, t*) or invalid (*h*′, *r*′, *t*′).

#### 2) Architecture

As mentioned above, *k* dimensional vectors of triple (*v*_*h*_, *v*_*r*_, *v*_*t*_) are considered as a matrix 𝒜 = [*v*_*h*_, *v*_*r*_, *v*_*t*_] ∈ ℝ^*k*×3^. For each 𝒜_*i*,:_ ∈ ℝ^*k*×3^ considered as row *i*^*th*^ of the matrix 𝒜.

Assuming that we use a filter *ω* ∈ ℝ^1×3^ in the convolutional layer, *ω* does not only verify the relationship between corresponding *v*_*h*_, *v*_*r*_, *v*_*t*_, but also generalizes its characteristics.

The filter *ω* slides on each row of matrix 𝒜 to create feature maps *v* = [*v*_1_, *v*_2_, …, *v*_*k*_] ∈ ℝ^*k*^:

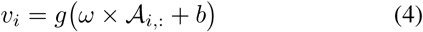

Where *b* ∈ ℝ is the bias and *g* is an activation function such as ReLU, sigmoid.

ConvKB uses different filters ∈ ℝ^1×3^ to create feature maps. The set of filters marked as Ω, and *τ* is marked as number of filters, such that: *τ* = |Ω | and have *τ* feature maps. These feature maps *τ* are concatenated to form a feature vector ∈ ℝ^*τk*×1^, then is multiplied with weight vectors **w** ∈ ℝ^*τk*×1^ and returns a score for a triple (*h, r, t*).

We defined the score function *f* of ConvKB following this equation:

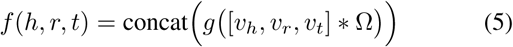

Where Ω, **w** are the shared independent parameters of *h, r, t*; is convolution operator and *concat* is the concatenation function of vector.

If we use one filter *ω, τ* = 1 and bias *b* = 0 with the activation function *g*(*x*) = |*x*| or *g*(*x*) = *x*^2^, and *ω* = [1, 1, 1] and **w** = [1, 1, …1] ∈ ℝ^*k*×1^. During the training phase, the ConvKB model becomes TransE. Thus, we can consider that TransE (or more generally TransH, TransD, TransR) is a special case of ConvKB. The Fig. 1 describe the computational process of the ConvKB model.

**Fig. 1.**
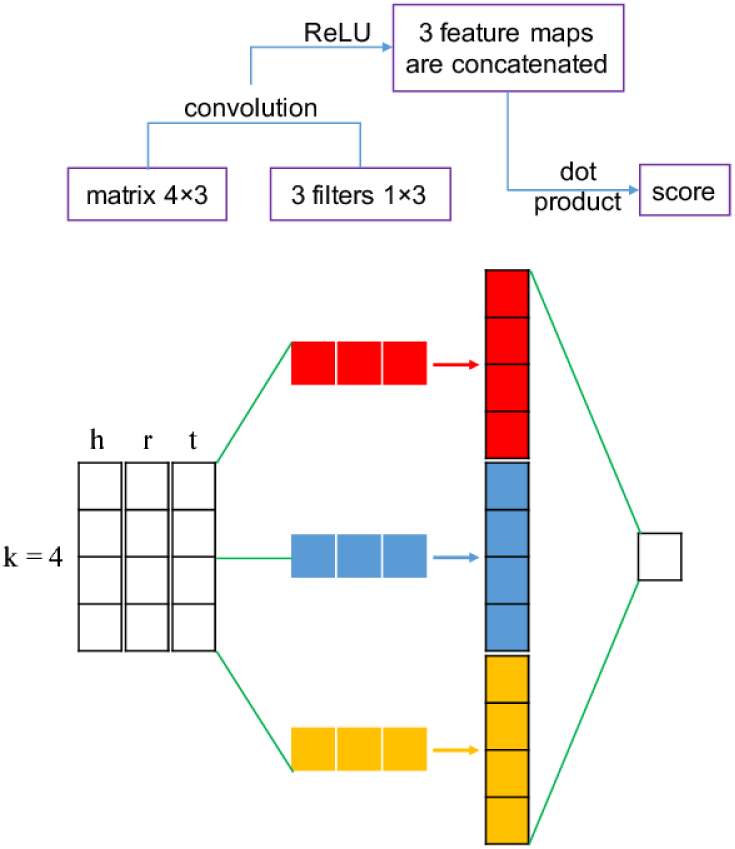
ConvKB Model [31].

#### 3) Loss Function

The ConvKB model uses the ADAM optimizer [33] during the training phase to optimize the loss function ℒ with *L*_2_ regularization over the weight **w**:

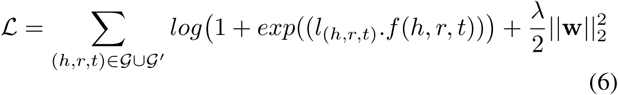

Where:

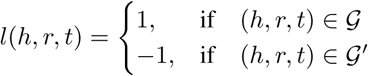

𝒢′ is a set of generated triples which are created from the original set of triples 𝒢.

#### 4) Applying ConvKB model for OsGenePrio dataset

Because they are based on vertices and edges structure, knowledge graphs have a wide benefits. In our model, rice genes and their attributes are represented by a set of triplets. In principle, each triplet is a set of 3 elements: head, relation and tail (*h, r, t*). Head stands for a gene id, tail stands for a gene attribute and relation stands for a relationship between gene id and its attribute extracted from different databases. Then, we consider vertex as head and tail of the triplet while edge is its relation. Currently, *OsGenePrio* contains only the relationship between genes and their attributes. The direct relation between genes is not available but it can be stated based on the same attributes belonging in the same cluster of genes. In section III-B2 and section III-B3, we present how vectors of gene id *v*_*h*_ are used to cluster and find similar genes.

## III. Experiments

### A. Dataset

To explore the information, we use PyRice package [16] to collect gene information from six databases: Oryzabase, RAP-DB, Gramene, Funricegene-genekeywords, Funricegene-faminfo, Funricegene-geneinfo.

Moreover, using the Uniprot and PRIN databases, we added more data for the training set. Due to the large number of gene attributes, we kept only some specific information taking into account a broader covering knowledge and a minimum of redundancy. In the following are presented the attributes used for our embedding model:

1. Protein_Uniprot;
2. GO_Uniprot or PyRice;
3. Keyword_Uniprot;
4. Description_PRIN;
5. Location_PRIN;
6. Position_PyRice;
7. CGSNL Gene Name_PyRice;
8. Trait Class_PyRice;
9. TO_PyRice;
10. PO_PyRice;
11. Keyword_Pyrice;
12. Name_PyRice.

#### 1) Data standardization

The tuning process to standardize the dataset took a lot of time to complete. After implement statistics of gene attributes, we found that attributes which have a high frequency of occurrence such as KW-0181, KW-1185 are keyword attributes specific from the Uniprot database and did not generalize for genes databases. Besides, some other attributes appear rarely, such as some annotations with gene ontology (GO), trait ontology (TO) and plant ontology (PO) attributes. These low frequency attributes may affect the training process. Therefore, the selection of attributes is also considered as a tuning parameter. This selection is very important for the quality of the dataset. After evaluating the dataset attributes, we removed those with the number of occurrences higher than 7000 and less than two. For instance, from Fig. 2 we can see that there are approximately 4000 attributes appear 1-5 times.

**Fig. 2.**
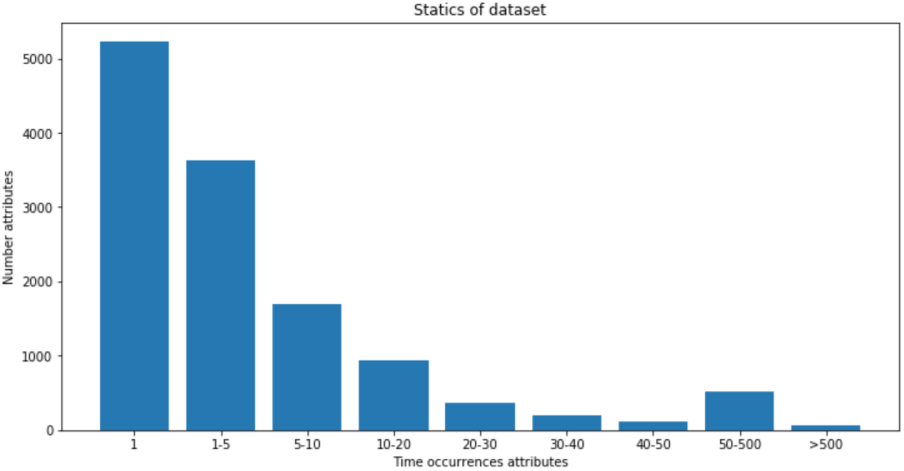
Statics dataset.

With gene attributes, some of them will be considered as a group. Here are some examples:

*26S proteasome regulatory subunit 7B*

*26S proteasome regulatory subunit RPN7*

…

*ABC transporter G family member 32*

*ABC transporter G family member 35*

…

*Chloroplastic group IIA intron splicing facilitator CRS1*

*Chloroplastic group IIB intron splicing facilitator CRS1*

…

Here, the issue is how to assign different properties to each group. Solution: using the jaccard similarity measurement with the following steps:

- Split attributes by length.
- Use the Jaccard distance equation: the number of overlap word based on 2 attributes A,B divide by the mean of 2 attributes.

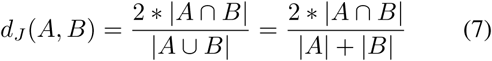 |*A* ∩ *B*|: number of overlap word at the same position in A and B. |*A*|, |*B*|: A and B are length of attributes.
- Two attributes will be considered in the same group if its distance is bigger than 0.6. Through experiments, we found that this value is suitable to fit with the dataset.

#### 2) Dataset for embedding model

The number of entities includes: 12 relations between entities (#R), 33268 entities (#E); where:

- 25742 is the number of gene identifiers;
- 7525 is the number of gene attributes on different databases

Table I show the statistics of the dataset OsGenePrio. The validation set is used to evaluate and optimize the model during the training phase. The number of triple after extraction is 206795 which is split into training and validation set.

**TABLE I.** Description dataset.

| Dataset | #R | #E | Triples in train | Triples in valid |
| --- | --- | --- | --- | --- |
| OsGenePrio | 12 | 33268 | 196455 | 10340 |

### B. Evaluation model

#### 1) Predict entities with triples

To predict the entity present at the head or the tail, for instance, to predict *h* given that (?, *r, t*) or predict *t* given that (?, *r, t*); with ? is the entity needed to be predicted. The output will be evaluated by ranking the score after applying *f* (*h, r, t*) with the validation set.

Each triple (*h, r, t*) is randomly replaced by a set of the first or last entity (candidate set). Then it is ranked by score (raw evaluation). Besides, there is a simplified approach used to “Filtered” setting protocol [25] which is similar to raw filter, but the triple is generated randomly from the first and last entity. If they exist in the training set, the triple will be removed and not be ranked during the evaluation process. We used three metrics to evaluate the ranking: Mean rank (MR), Mean reciprocal rank (MRR) and Hit ratio (Hits@).

#### 2) K-Means to cluster genes

From ConvKB results, we obtained a vector of genes to cluster and classify in order to identify similar genes. Alternatively, we can predict attributes and values of a gene. However, the predictions need to be confirmed by biologists, thus this part will not be evaluated in the scope of this paper.

To evaluate the K-Means clustering, we used the total distance from genes in the cluster to the centroid of the cluster. From this cluster, we can capture the general characteristic about a group of genes. For example, this cluster will represent a trait or a disease.

#### 3) K-nearest neighbors to find similarity of gene

Find K similar genes corresponding to a gene input or rank genes based on the similarity of attributes. The score will be high if two genes are similar. This work is very potential in term of improving time and performance.

### C. Training model

Based on the research of Wang et al. [34], [35], we applied the Bernoulli’s distribution to generate the head and tail of sampling invalid triples. As we mentioned in the previous part, TransE is used as the pre-trained model to initialize embedding entities while ConvKB is used as pre-trained model to embed relationships among entities. During the training phase, the details of parameters for TransE are: the number of epochs 120, the dimensions of embeddings *k* ∈ {50, 100, 150}, ADAM learning rate *η* ∈ {1e^−3^, 1e^−4^}, *L*_2_ norm and margin *γ* ∈ {1, 3, 5}. To fit the model parameters including the entity and relation to the dataset, filters *ω* and the weight vector **w**, we choose the initial learning rate *η* ∈ {1e^−3^, 1e^−4^} with ADAM optimizer and ReLU as activation function *g*. Batch size is set at 128 with the *L*_2_ regularizer *λ* = 1e^−3^. The filters *ω* are initialized by a truncated normal distribution or by [0.1, 0.1, − 0.1] with number of filters *τ* ∈ {100, 150, 200}. To train ConvKB, the number of epochs is set up to 125. The output from last epochs is used for the evaluation later.

### D. Main experimental results

In this section, we will use different set of parameters to evaluate and select the best ones for the model. Parameters are selected based on the results obtained on the validation set.

#### 1) Link prediction results

To ensure about the results, we evaluated the model based on 3 metrics which are proposed above MR, MRR and Hits@. We selected the best set of parameters to compare with MRR. Results are presented below. The highest MMR scores on the validation set are obtained when using *k* = 150, *τ* = 150, learning rate *η* = 1e^−3^ (for both TransE and ConvKB), margin *γ* = 3, the truncated normal distribution for filter initialization. Table II shows the results based on the validation set with the configuration “Filtered” and “Not Filtered” protocol.

**TABLE II.**
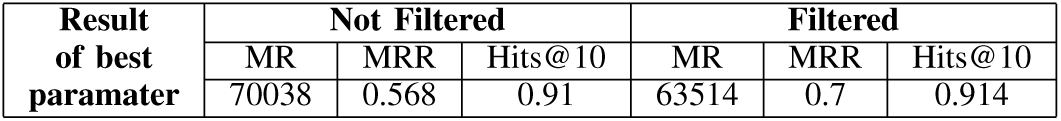
Results on the validation set with the best set of parameters.

| Result of best parameter | Not Filtered |  |  | Filtered |  |  |
| --- | --- | --- | --- | --- | --- | --- |
|  | MR | MRR | Hits@10 | MR | MRR | Hits@10 |
|  | 70038 | 0.568 | 0.91 | 63514 | 0.7 | 0.914 |

#### 2) K-Means results

Fig. 3 shows that the total distance decreases when we increase the number of clusters. We can see with cluster higher 300, the loss of K-means start the convergence. The number of clusters recommended from data scientists is around *k* = [75, 100] clusters. Moreover, following Fig. 3, we can see that the number of recommended clusters is suitable.

**Fig. 3.**
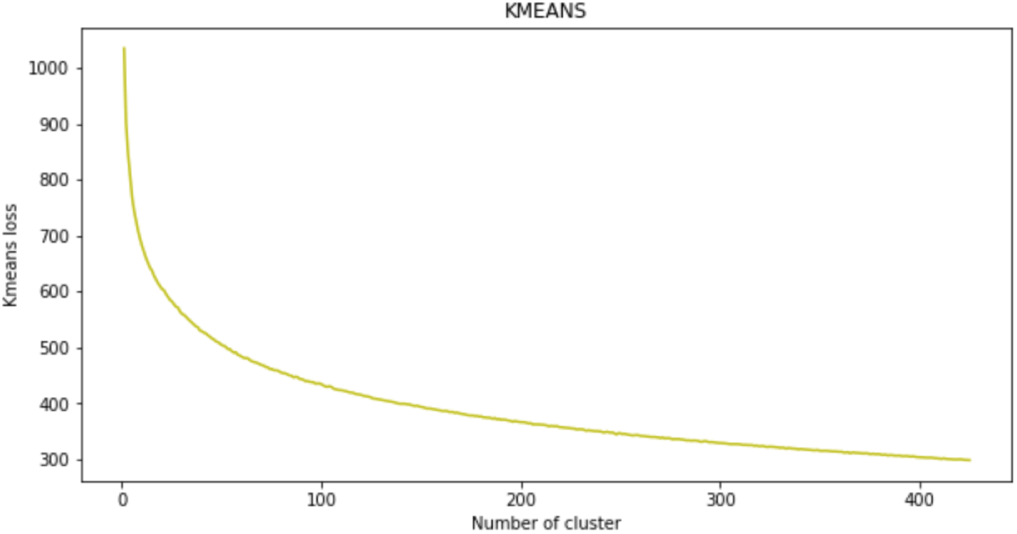
Performance on K-Means.

#### 3) K-nearest neighbors results

From the Fig. 4 we can see a lot of similar attributes between the gene search and the proposed gene model.

**Fig. 4.**
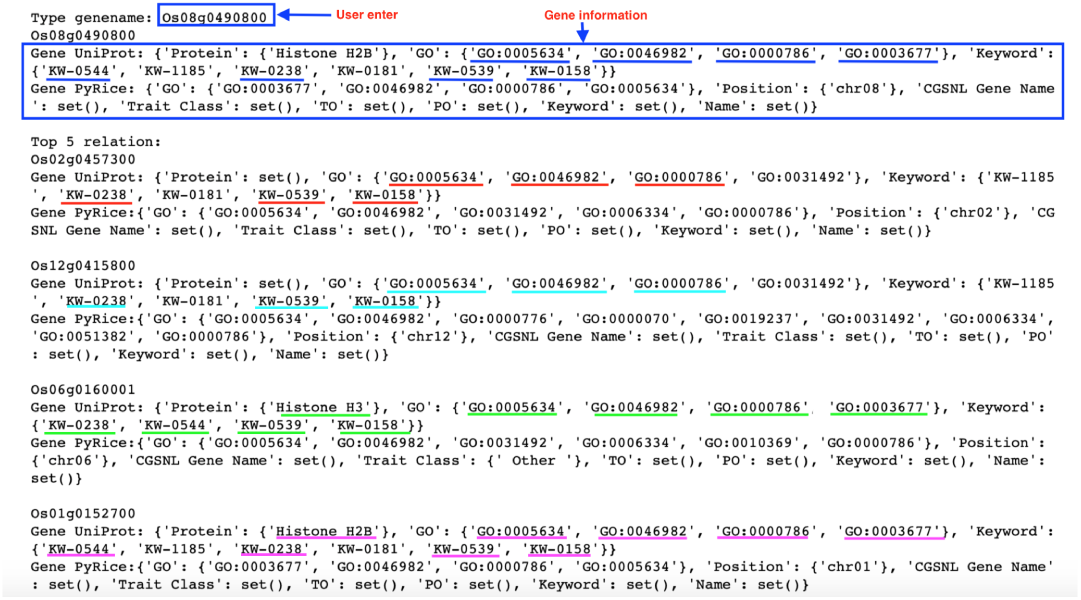
Example of using K-nearest neighbor.

## IV. Future work

Through the evaluation of experimental results on the dataset, we also realized the drawback of the ConvKB model.

In the future, we aim to improve the results of the model following the directions bellow:

- Improved the PyRice tool: integrating the ConvKB model will make gene search more efficient and meaningful, not just based on regular text searches. Besides, we can expand to other databases on rice genes.
- Improvements in data: since there is still some noise, for example, the gene attribute groups are not fully homogeneous. This issue needs the attention of researchers in biology as well as additional methods to a better pre-processing phase. Besides, the important gene information might be missing, which leads to difficulties in training and prediction.
- Model enhancements: the new model only uses TransE, which is the simplest models in the knowledge graph and CNNs. However, recently, other models have been developed to improve the performance in terms of structure and efficiency.

## V. Conclusion

This project focused on the studies of knowledge graphs combining with CNNs to apply on gene analysis. The proposed method implemented from a new dataset OsGenePrio which is built from publishing databases of plant science. The result shows that CNN is very exploitable in other fields, not only in computer vision. Besides, it also shows the advantages compared to traditional embedding methods such as translating embedding model.

To sum up, the project has completed the proposed goals in the objective. First, we developed some tools and generated a dataset for genetic analysis, then do some research based on knowledge graph and set up some tests and evaluation methods. Finally, we provided an evaluation of the results.

## Acknowledgment

The authors would like to thank IRD and ICTLab for their supports.

## References

[1] Y. Moreau and L. C. Tranchevent, “Computational tools for prioritizing candidate genes: Boosting disease gene discovery,” Nature Reviews Genetics, vol. 13, no. 8, pp. 523–536, 2012.

[2] L. Hou and H. Zhao, “A review of post-GWAS prioritization approaches,” Frontiers in Genetics, vol. 4, no. DEC, pp. 2009–2014, 2013, iSBN: 1664-8021 (Print) r1664-8021 (Linking).

[3] L. C. Tranchevent, A. Ardeshirdavani, S. ElShal, D. Alcaide, J. Aerts, D. Auboeuf, and Y. Moreau, “Candidate gene prioritization with Endeavour,” Nucleic acids research, vol. 44, no. W1, pp. W117–W121, 2016.

[4] A. A. Kumar, L. Van Laer, M. Alaerts, A. Ardeshirdavani, Y. Moreau, K. Laukens, B. Loeys, and G. Vandeweyer, “pBRIT: gene prioritization by correlating functional and phenotypic annotations through integrative data fusion,” Bioinformatics (Oxford, England), vol. 34, no. 13, pp. 2254–2262, Jul. 2018.

[5] M. Pérez-Enciso and L. M. Zingaretti, “A guide on deep learning for complex trait genomic prediction,” Genes, vol. 10, no. 7, 2019.

[6] A. Rao, S. Vg, T. Joseph, S. Kotte, N. Sivadasan, and R. Srinivasan, “Phenotype-driven gene prioritization for rare diseases using graph convolution on heterogeneous networks,” BMC medical genomics, vol. 11, no. 1, pp. 57–57, Jul. 2018. [Online]. Available: https://www.ncbi.nlm.nih.gov/pubmed/29980210

[7] N. Kurata and Y. Yamazaki, “Oryzabase. an integrated biological and genome information database for rice,” Plant physiology, vol. 140, no. 1, pp. 12–17, 2006.

[8] H. Sakai, S. S. Lee, T. Tanaka, H. Numa, J. Kim, Y. Kawahara, H. Wakimoto, C.-c. Yang, M. Iwamoto, T. Abe et al., “Rice annotation project database (rap-db): an integrative and interactive database for rice genomics,” Plant and Cell Physiology, vol. 54, no. 2, pp. e6–e6, 2013.

[9] L. Mansueto, R. R. Fuentes, F. N. Borja, J. Detras, J. M. Abriol-Santos, D. Chebotarov, M. Sanciangco, K. Palis, D. Copetti, A. Poliakov et al., “Rice snp-seek database update: new snps, indels, and queries,” Nucleic acids research, vol. 45, no. D1, pp. D1075–D1081, 2016.

[10] W. Yao, G. Li, Y. Yu, and Y. Ouyang, “funricegenes dataset for comprehensive understanding and application of rice functional genes,” Gigascience, vol. 7, no. 1, p. gix119, 2017.

[11] U. Consortium, “Uniprot: a worldwide hub of protein knowledge,” Nucleic acids research, vol. 47, no. D1, pp. D506–D515, 2018.

[12] M. K. Tello-Ruiz, S. Naithani, J. C. Stein, P. Gupta, M. Campbell, A. Olson, S. Wei, J. Preece, M. J. Geniza, Y. Jiao et al., “Gramene 2018: unifying comparative genomics and pathway resources for plant research,” Nucleic acids research, vol. 46, no. D1, pp. D1181–D1189, 2017.

[13] I. P. Consortium, “Information commons for rice (ic4r),” Nucleic acids research, vol. 44, no. D1, pp. D1172–D1180, 2015.

[14] Y. Kawahara, M. de la Bastide, J. P. Hamilton, H. Kanamori, W. R. McCombie, S. Ouyang, D. C. Schwartz, T. Tanaka, J. Wu, S. Zhou et al., “Improvement of the oryza sativa nipponbare reference genome using next generation sequence and optical map data,” Rice, vol. 6, no. 1, p. 4, 2013.

[15] H. Gu, P. Zhu, Y. Jiao, Y. Meng, and M. Chen, “Prin: a predicted rice interactome network,” BMC bioinformatics, vol. 12, no. 1, p. 161, 2011.

[16] Q. Do, P. Larmande, and H. B. Hai, “Pyrice: a python package for querying oryza sativa functional databases,” unpublished.

[17] G. Kasneci, F. M. Suchanek, G. Ifrim, M. Ramanath, and G. Weikum, “Naga: Searching and ranking knowledge,” in 2008 IEEE 24th International Conference on Data Engineering. IEEE, 2008, pp. 953–962.

[18] C. Xiong, R. Power, and J. Callan, “Explicit semantic ranking for academic search via knowledge graph embedding,” in Proceedings of the 26th international conference on world wide web. International World Wide Web Conferences Steering Committee, 2017, pp. 1271–1279.

[19] Y. Hao, Y. Zhang, K. Liu, S. He, Z. Liu, H. Wu, and J. Zhao, “An end-to-end model for question answering over knowledge base with cross-attention combining global knowledge,” in Proceedings of the 55th Annual Meeting of the Association for Computational Linguistics (Volume 1: Long Papers), 2017, pp. 221–231.

[20] B. Yang and T. Mitchell, “Leveraging knowledge bases in lstms for improving machine reading,” in Proceedings of the 55th Annual Meeting of the Association for Computational Linguistics (Volume 1: Long Papers), 2017, pp. 1436–1446.

[21] R. West, E. Gabrilovich, K. Murphy, S. Sun, R. Gupta, and D. Lin, “Knowledge base completion via search-based question answering,” in Proceedings of the 23rd international conference on World wide web. ACM, 2014, pp. 515–526.

[22] A. Bordes, J. Weston, R. Collobert, and Y. Bengio, “Learning structured embeddings of knowledge bases,” in Twenty-Fifth AAAI Conference on Artificial Intelligence, 2011.

[23] M. Nickel, K. Murphy, V. Tresp, and E. Gabrilovich, “A review of relational machine learning for knowledge graphs,” Proceedings of the IEEE, vol. 104, no. 1, pp. 11–33, 2015.

[24] B. Yang, W.-t. Yih, X. He, J. Gao, and L. Deng, “Embedding entities and relations for learning and inference in knowledge bases,” arXiv preprint 1412.6575, 2014.

[25] A. Bordes, N. Usunier, A. Garcia-Duran, J. Weston, and O. Yakhnenko, “Translating embeddings for modeling multi-relational data,” in Advances in neural information processing systems, 2013, pp. 2787–2795.

[26] T. Trouillon, J. Welbl, S. Riedel, É. Gaussier, and G. Bouchard, “Complex embeddings for simple link prediction,” in International Conference on Machine Learning, 2016, pp. 2071–2080.

[27] Y. LeCun, L. Bottou, Y. Bengio, P. Haffner et al., “Gradient-based learning applied to document recognition,” Proceedings of the IEEE, vol. 86, no. 11, pp. 2278–2324, 1998.

[28] R. Collobert, J. Weston, L. Bottou, M. Karlen, K. Kavukcuoglu, and P. Kuksa, “Natural language processing (almost) from scratch,” Journal of machine learning research, vol. 12, no. Aug, pp. 2493–2537, 2011.

[29] Y. Kim, “Convolutional neural networks for sentence classification,” in Proceedings of the 2014 Conference on Empirical Methods in Natural Language Processing (EMNLP), 2014, pp. 1746–1751.

[30] T. Dettmers, P. Minervini, P. Stenetorp, and S. Riedel, “Convolutional 2d knowledge graph embeddings,” in Thirty-Second AAAI Conference on Artificial Intelligence, 2018.

[31] D. Q. Nguyen, T. D. Nguyen, D. Q. Nguyen, and D. Phung, “A novel embedding model for knowledge base completion based on convolutional neural network,” in Proceedings of the 2018 Conference of the North American Chapter of the Association for Computational Linguistics: Human Language Technologies, Volume 2 (Short Papers), 2018, pp. 327–333.

[32] D. Q. Nguyen, T. Vu, T. D. Nguyen, D. Q. Nguyen, and D. Phung, “A capsule network-based embedding model for knowledge graph completion and search personalization,” in Proceedings of the 2019 Conference of the North American Chapter of the Association for Computational Linguistics: Human Language Technologies, Volume 1 (Long and Short Papers), 2019, pp. 2180–2189.

[33] D. P. Kingma and J. Ba, “Adam: A method for stochastic optimization,” arXiv preprint 1412.6980, 2014.

[34] Z. Wang, J. Zhang, J. Feng, and Z. Chen, “Knowledge graph embedding by translating on hyperplanes,” in Twenty-Eighth AAAI conference on artificial intelligence, 2014.

[35] Y. Lin, Z. Liu, M. Sun, Y. Liu, and X. Zhu, “Learning entity and relation embeddings for knowledge graph completion,” in Twenty-ninth AAAI conference on artificial intelligence, 2015.

